# Behaviour and cold hardiness of the purple stem borer in winter, colonizing more northerly latitudes

**DOI:** 10.1101/819417

**Authors:** Jianrong Huang, Guoping Li, Haixia Lei, Chunbin Fan, Caihong Tian, Qi Chen, Bo Huang, Huilong Li, Zhaocheng Lu, Hongqiang Feng

## Abstract

To escape or alleviate low temperatures in winter, insects have evolved many behavioral and physiological strategies. The rice pest insect, the purple stem borer, *Sesamia inferens* (Walker) is currently reported to be expanding their northern distributions and causing damage to summer maize in Xinxiang, China. However, their method of coping with the lower temperature in the new northern breeding area in winter is largely unknown. This paper investigates the overwinter site of *S. inferens*, and identifies the cold hardiness of larvae collected from a new breeding area in winter and explores a potential distribution based on low temperature threshold and on species distribution model, MaxEnt. The results show that the overwintering location of the *S. inferens* population is more likely to be underground with increasing latitude and,in the north, with the temperature decreasing, the larvae gradually moved down the corn stalk and drilled completely underground by February 18^th^. Those who were still above ground were all winterkilled. The cold hardiness test shows the species is a moderate freeze-tolerant one, and Supercooling Points (SCP), Freezing Points (FP) and mortality rate during the middle of winter (January, SCP: −7.653, FP: −6.596) were significantly lower than early winter (October) or late winter (March). Distribution in the new expansion area was predicted and the survival probability area was below N 35° for the Air Lower Lethal Temperature (ALLT_50_) and below N 40° for the Underground Lower Lethal Temperature (ULLT_50_), The suitable habitat areas for *S. inferens* with MaxEnt were also below N 40°. This study suggests the overwinter strategies have led to the colonization of up to a five degree more northerly overwintering latitude. This behavior of *S. inferens* could help maize producers to propose a control method to increase pest mortality by extracting the maize stubble after harvest.

## 1 Introduction

Temperature is the main abiotic factor that determines the growth and breeding of ectotherms. The distribution of most insects is directly related to extreme temperature (Sunday et al., 2012). The minimum temperature in winter which determines the survival rate of insect wintering populations is an important factor limiting the potential geographical distribution of insects (Bale, 1996; Stahl et al., 2006), particularly in ectotherms, setting northern range limits (Stahl et al., 2006; Gray, 2008; Calosi et al., 2010). The low temperature in winter constrains the behavioral strategies of colonial insects (Labrie et al., 2008), such as migration, drilling holes in refuges or making a thick cocoon, to avoid winterkilling. Insects also avoid chill injury by regulating physiological metabolism and accumulation of cold-tolerant substances. The warming of the global climate also leads to poleward distribution in some species’ ranges, and may result in insects evolving new overwintering strategies in different breeding areas (Reynolds et al., 2015; Hu et al., 2015).

The purple stem borer, *Seramia inferens* Walker, belongs to the order Lepidoptera, Noctuidae, and *Sesamia*. They are polyphagous insects and their hosts are mainly gramineous crops or weeds such as rice, wheat, maize and they could live for a whole generation on rice, water shoots, and maize (Guo, 2017). With the changes in the farming system, crop distribution, and use of pesticides, *S. inferens* is gradually becoming a significant rice pest in many parts of China (Chen et al., 2015). *S. inferens* is distributed in rice-producing countries, mainly in Asia, the India Peninsula, Bangladesh, Myanmar, Thailand, Malaysia, Taiwan, Pakistan and Japan. In China, the species is distributed south of N 34 degrees latitude (Zhang, 1965), The ranges are consistent with research by Ezcurra et al., (1978). Based on the cold resistant characteristics and field experiments of *S. inferens*, the distribution area was also agreed to be south of N 34 degrees latitude by Gu (1985). However, adult *S. inferens* were first found by light trap and also caused maize plant damage in Xinxiang City, Henan Province, China (E: 113.696°, N: 35.021°) in 2014 (Huang and Feng, 2015). Further investigation illustrated that *S. inferens* can produce three generations in total and overwinter in local areas, following field surveys (Huang et al., 2017). They must have developed tolerance to cold to establish a population in the new northern breeding area in China in winter. Past research mainly focuses on overwintering in rice stubble in southern areas. It states that *S. inferens* overwinter as late instar larvae, and it is an exclusively freeze-tolerant insect (Guo et al., 2002; Sun et al., 2014). Significant damage to maize crops has also been reported in southern China (Xie et al., 1985). However, studies on overwintering strategies of the pest in the maize planting area and cold tolerance of *S. inferens* in the new expansion area remain inadequate.

In this study, the behavioral strategies were confirmed by investigating the overwinter site and survival of *S. inferens* in early-middle-late winter in the field. The diapause larvae were also collected and the cold hardiness was measured, and a regional distribution based on low temperature threshold and species distribution model MaxEnt were used to generate a potential distribution of *S. inferens*. This research aims to identify the methods of *S. inferens* larvae to tolerate cold temperatures and to study the northern distribution boundary in the expansion zone.

## 2 Materials and Methods

### 2.1 Latitudinal overwintering position of *S. inferens*

Field surveys were carried out in the late overwinter period on the 5^th^ of February 2016 in Nanjing (NJ) (N 32.65°, E 119.43°) and the 5^th^ of February 2016 in Changsha (CS) (N 28.47°, E 113.35°) in remaining rice stubble. *S. inferens* larvae were also investigated on the 18^th^ of February 2016 in Xinxiang (XX) (N 35.02°, E 113.69°) in remaining maize stalks. The number of *S. inferens* at each location on the rice plants was recorded. Four vertical positions of the remaining rice or maize stalk from top to bottom were devised, including >20 cm above ground (AG), 10-20 cm AG, 0-10 cm AG and underground. The data of Xianyou (XY) (N 25.36°, E 118.69) samples cited in Gu (2014) were documented in a field survey on the 2^nd^ and 11^th^ of February in 2013.

### 2.2 Underground-forward behavior on corn stalks

The vertical distribution of *S. inferens* larvae in maize plants was destructively sampled to determine survival of field conditions during the winters of 2015-2016 on the 6^th^ of October, 2^nd^ of November, 10^th^ of December and 18^th^ of February in XX. Numbers of living and dead *S. inferens* larvae and the position of those insects in the maize stalk were recorded. The mortality ratio at each location on the maize plant was calculated. Six vertical positions of the maize stalk, which remained completely in the field except the corn ear, from top to bottom are defined as positions 1-6. Position 1 represents the high maize ear section. Position 2 represents two sections of the maize ear. Position 3 represents two sections under the maize ear. Position 4 represents three sections above the soil surface. Position 5 represents the base of the maize stem and maize root under the soil surface. Position 6 represents the surrounding soil near the root.

### 2.3 Cold hardiness during winter

Wild *S. inferens* larvae were collected on remaining summer maize stalks in Xinxiang City (N 32°, E 119°) in different winter periods on October 31^st^ and November 10^th^ in 2015, January 27^th^, February 18^th^ and March 30^th^ in 2016. Each larva replicate was placed in a 0.5 ml plastic tube with a 1-2 mm diameter ventilation hole in the lid and labeled. A weight test was then carried out (Mettler Toledo ME204, Zurich, Switzerland). The Supercooling Points (SCP) and Freezing Points (FP) for each larval replicate were then measured by a small thermocouple thermometer (Liu et al., 2016) (Temp 20, Beijing Pengcheng Electronic Technology Center), which meant placing the thermometer into the plastic tube and touching the larvae bodies immobilized with cotton wool. Thermocouples were attached to a multichannel data logger (USB-TC, Measurement Computing, Norton, MA), and recorded and logged by using Tracer-DAQ software (Measurement Computing). The tubes were then placed in a −20°C freezer for 30 minutes. The body temperature of the larvae was gradually reduced, so that all larval replicates could have SCP and FP recorded, as in Leather (1993). After testing, death was assessed by the lack of mandibular and body movement after 6 hours recovery at 24°C and a mortality rate was identified by the chill-coma recovery numbers. All sample larvae were then dried at 65°C for 48 h in a drying oven and then weighed again, to determine the water content of each larva.

### 2.4 Lethal low temperature experiment

Larvae were collected in Anyang (N 35.57, E 114.85) in the same field on the 5^th^ of November 2015 and each larva was also placed in a separate 0.5 ml plastic tube as mentioned above. 20 cm of soil was placed in a container (L605×W410×H240 mm), along with moist cotton wool at the bottom of the container in order to guarantee the same humidity as the overwintering period. The tubes with larvae were then placed vertically into soil in the container. 2 cm of soil was placed on the tube to simulate the overwintering environment of *S. inferens* underground. Because the larvae were diapaused, there was no need for food to survive. The container was placed in an outdoor environment. Two microclimate sensors (TH11R, HHW) were installed in air (10 cm from the soil surface) with a shelf, and at 2 cm soil depth to record the temperature of both sites until March 30^th^ 2016. Low-temperature exposure experiments were carried out to measure the thresholds for the long-term survival of *S. inferens* at a constant temperature, separated into 4 times (Nov 9^th^, Dec11^th^, Jan 28^th^, Feb 19^th^) and 7 gradient low temperatures (0°C, −5°C, −10°C, −15°C, −20°C, −25°C, and −30°C). 7×20 larval replicates were treated each time and each larva was also placed in a separate 0.5 ml plastic tube as mentioned above. All replicates were placed in a 0°C freezer for a 12 hour cold treatment, and then 20 replicates were taken out and placed at ambient temperature for 24 hours to determine whether they were alive when moving or dead. The number of dead larvae was recorded. Those remaining replicates then experienced −5 °C for 12 hours by reducing the freezer to the next temperature (−10 °C), and then 20 replicates were taken out to check if alive or dead, and so forth until all 7×20 larval replicates were tested. Ultimately, the mortality rate of different temperatures was identified. A low temperature threshold: Air Lower Lethal Temperature (ALLT_50_), which is defined as the temperature where 50% of individuals die in the low-temperature exposure experiments mentioned above, was calculated by a logistic linear regression (Andersen et al., 2015). In this study, because the sampled larvae on different dates showed similar mortality rates after exposure to different temperatures, the data of the sample dates were taken together to calculate the ALLT_50,_ and the ALLT_50_ value was −4.5 °C.

### 2.5 The distribution based on low temperature threshold

Daily minimum temperatures in 47 different cities (Henan province, Hebei province, Tianjing city and Beijing city were included, and those cities formed a south-north belt in north plain of China, in order to illustrate the northern overwinter site of *S. inferens*) from 2007 to 2014 were collected from national weather stations (https://data.cma.cn/). Previous research shows the minimum temperatures above ground were linearly correlated with minimum temperatures underground (Parton and Logan, 1981; Horton, 2012). In this study, the winter temperatures from November to March were recorded in the air (10 cm from the soil surface) and underground (at 2 cm soil depth) by two microclimate sensors mentioned above and it was found that there is an obvious significant linear relationship between lowest daily air and lowest underground temperature (*R*^2^=0.9161, *p*<0.001, *n*=100). Based on this linear model, the minimum daily air temperatures in different cities were translated into the minimum daily soil temperature at 2 cm underground because the underground data were not available at national weather stations and the landscape and soil composites of the 47 cities were very similar. Another low temperature threshold, Underground Lower Lethal Temperature (ULLT_50_) was also translated by linear equation and the value was −9.7 °C, i.e. when the air temperature was −9.7 °C the soil temperature postulated reached −4.5 °C. 47 location survival rates of air and underground was speculated by predicting all larvae would die when the annual daily minimum temperature was lower than the ALLT_50_ or underground, respectively, i.e. if the annual minimum temperature is lower than the ALLT_50_ or ULLT_50_, 0% of the population will survival. 100% will survival if the annual minimum temperature is higher than the ALLT_50_ or ULLT_50._ The calculation of survival rates in different cities from 2007 to 2014, resulted in identification of the total proportion of insect survival over the 8 years. Based on the proportion of each cities, a potential distribution based on annual minimum temperature threshold could be produced with ArcMap (version 10.2, ESRI, Redlands, CA, USA) (Hong et al., 2014).

### 2.6 The distribution based on MaxEnt

In order to calibrate the distribution mentioned above and make up for the deficiency of the limited field-investigation scale, a species distribution model MaxEnt (version 3.3.3k) was used to generate a potential district-level distribution map of *S. inferens*. 19 bioclimatic data layers from the WorldClimte dataset were obtained (Fick and Hijmans, 2017; http://www.worldclim.org/). 147 district (cities) occurrences were obtained from published articles or websites, including the new northern breeding region in China (Zhang et al, unpublished, S1 Table 1 and S1 Table 2). 60% of total district (city) occurrences were used for model calibration (training data: 98 districts) and the remaining for model validation (test data: 49 districts). GoogleEarth was used to generate approximate coordinates. The area under the ROC (receiver operating characteristic) curve metric was used to evaluate the model performance (Phillips et al., 2006; Kumar et al., 2014) (S1 Fig 1). Current and potential distribution maps were generated using ArcMap and three arbitrary categories were defined as low (<0.15), medium (0.15-0.53) and high (>0.53) based on predicted habitat suitability.

### 2.7 Statistics

Data were tested for normal distribution using the Shapiro-Wilk test and normally distributed data was compared by an independent samples t-test and Pearson’s correction test.The differences between factors were evaluated using Tukey-Kramer groupings comparison in the least squares means and Non-normally distributed data were compared by nonparametric tests of group differences. The linear or nonlinear models were identified with p<0.05 for the significance, and all data were analyzed with the software program R 3.5.3 (R Core Team 2017).

## 3 Results

### 3.1 Geographical variation in overwinter position

The *S. inferens* population was closer to the ground with increasing latitude in late winter and 100% and 75% of overwinter individuals were underground in northern locations in XX and NJ, respectively. But in southern regions, CS and XY, they were mainly 0-10 cm AG (Fig 1).

**Fig 1.**
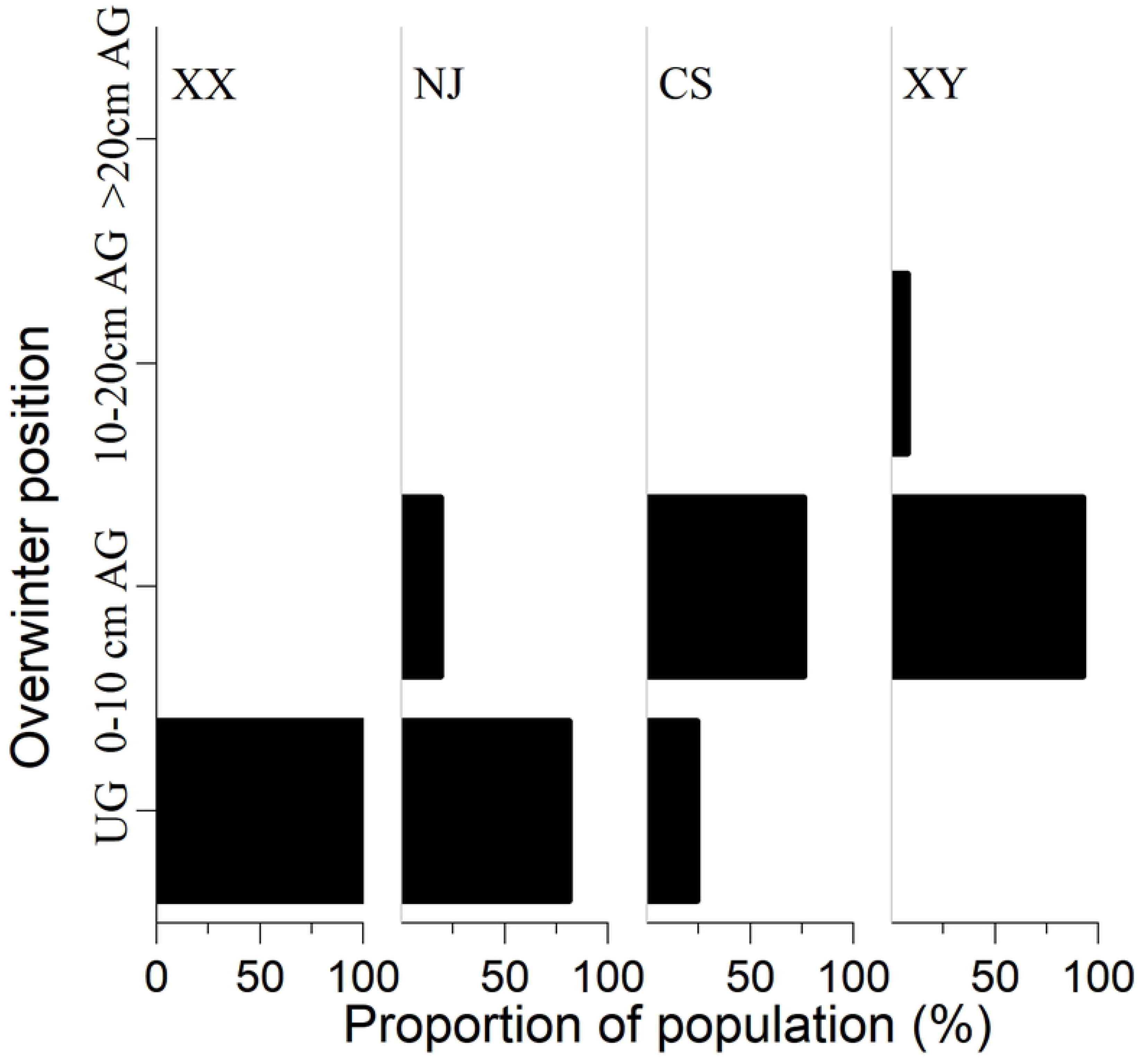
Geographical variation in overwintering position. UG: Underground; AG: above ground, XX: Xingxiang(*n*=19), NJ: Nanjing(*n*=74), CS: Changsha(*n*=33); XY: Xianyou(*n*=13).

### 3.2 Overwintering behavioral in maize stalk

Before overwintering, on the 6^th^ of October, 72.222% (n=18) of the living larvae were distributed into two sections of maize ear and two sections under the maize ear in the field (positions 2 and 3). No larvae were found on the underground part. On the 2^nd^ of November, the location of the population distribution shifted to the ground. 48.276% (*n*=29) of the population was located at positions 2 and 3, meanwhile, 13.793% of the population was on the underground part at the base of the maize stalk (position 5). On the 10^th^ of December, it was found that 75% (*n*=64) of the population was distributed in the underground part (position 5), and the number of insects at other locations was in total only 25%. On the 18^th^ of February, 100% (*n*=19) of the population had drilled into the underground position 5 (Fig 2).

**Fig 2.**
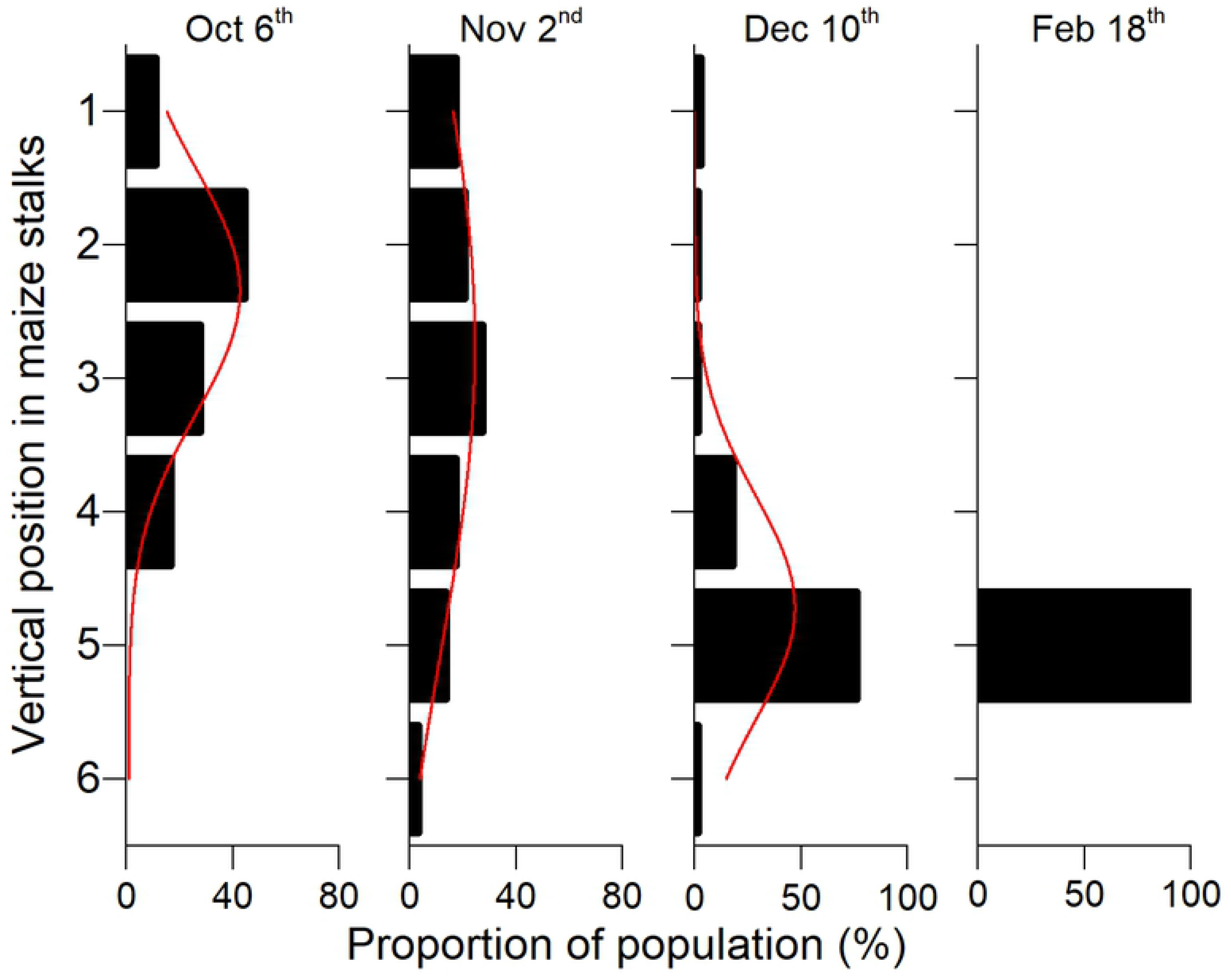
Underground-forward behavior of *S. inferens* in remaining summer corn stalks in winter.

The mortality rates which include above ground and underground *S. inferens* larvae were 25.5% (*n*=86) and 20.83% (*n*=25) on the 10^th^ of December and the 18^th^ of February in the field, respectively. 100% of total overwintering populations who remained AG in winter died, and only 1.802% of total overwintering populations who drilled into the ground died (Fig 3).

**Fig 3.**
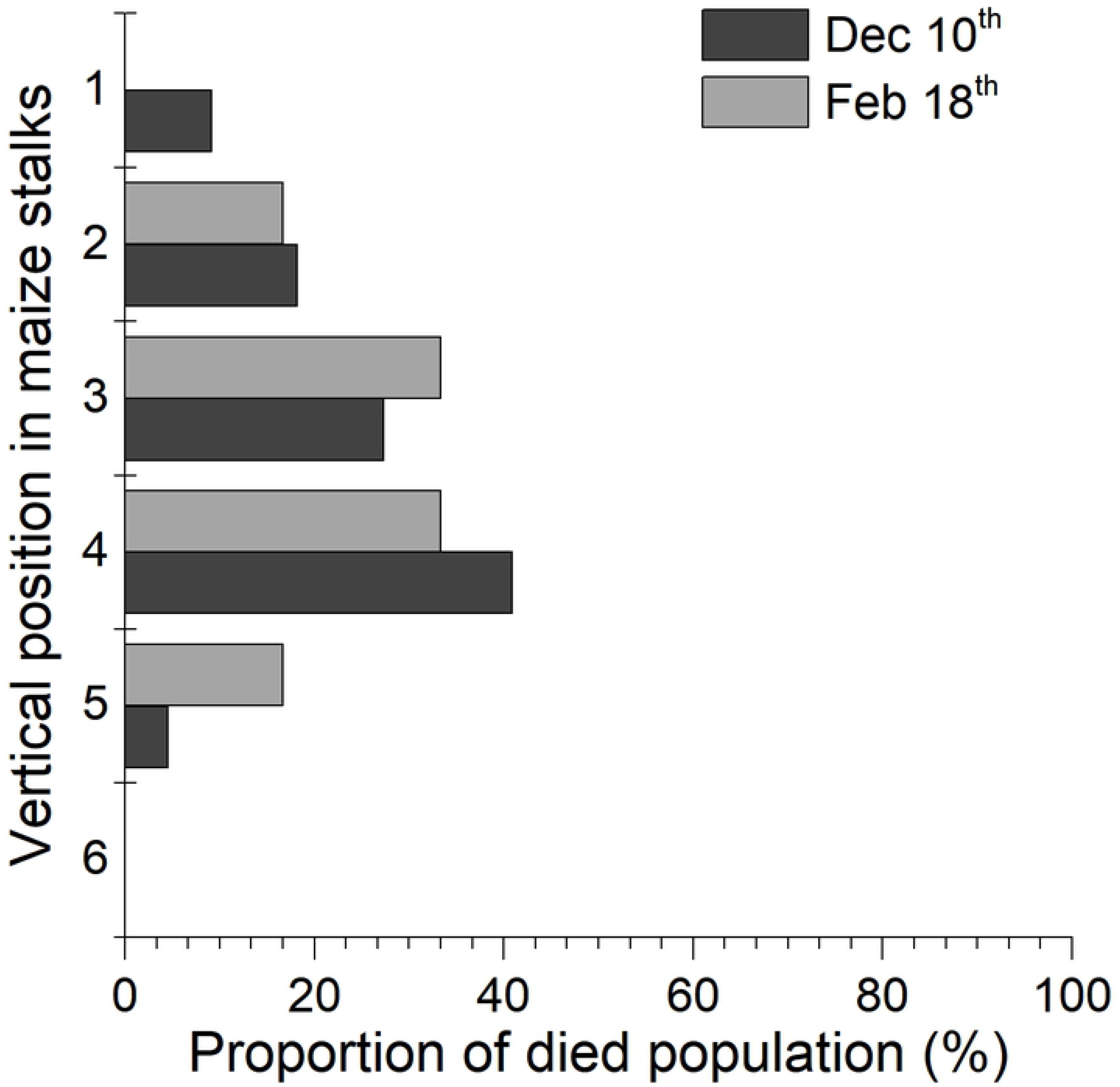
Proportion of *S. inferens* dead larvae in remaining winter maize stalks.

### 3.3 Cold hardiness of *S. inferens*

There were significant differences in SCP and FP of *S. inferens* larvae over time (SCP: *F*_4,91_=4.993, *p*= 0.003; FP: *F*_4,91_=10.26, *p*<0.001). The mean SCP (±SE) of *S. inferens* larvae ranged from the highest (−5.731±0.405 °C) in March to the lowest (−7.653±0.952 °C) in January (Fig 4) and the mean and minimum value of the SCP first decreased and then rose with the cold temperature event. The lowest value −20.02 °C appeared on the 27^th^ of Jan (Table 1). Similarly, the mean of the FP first decreased and then rose. The means of the FPs in middle winter (December: −5.49±0.447; January:−6.596±0.889) were significantly lower than early and end of winter (November, October and March). In contrast, there was no significant difference in the fresh weight of larvae during the cold temperature period (*F*_4,91_=1.997, *p*=0.102), but there was a significant gradual decrease in dry weight (*F*_4,91_=3.934, *p*=0.006) and water content of the larvae (*F*_4,87_=3.19, *p*=0.017). Larvae dry weight showed a decreased tendency throughout the winter and the water content decreased at the beginning of winter and slightly increased at the end of winter (Table 1).

**Table 1.** Minimum SCP, fresh mass, dry mass, water content and mortality rate of *S. inferens* larvae after SCP test.

| Date | *n* | Minimum SCP (°C) | Fresh mass (g) | Dry mass (g) | Water content (g) | Mortality rate after cold hardness test (%) | Daily mean air temperature (°C) |
| --- | --- | --- | --- | --- | --- | --- | --- |
| Oct 31<sup>st</sup> | 24 | -10.38 | 0.19±0.015 a | 0.069±0.007 ab | 0.121±0.008 a | 91.667 | 13.67 |
| Dec 10<sup>th</sup> | 20 | -11.95 | 0.179±0.019 a | 0.087±0.012 a | 0.092±0.01 ab | 30 | 2.18 |
| Jan 27<sup>th</sup> | 19 | -20.02 | 0.146±0.016 a | 0.051±0.007 bc | 0.094±0.01 ab | 15.789 | -3.24 |
| Feb 18<sup>th</sup> | 12 | -19.74 | 0.157±0.02 a | 0.069±0.011 ab | 0.088±0.01 b | 37.5 | 3.03 |
| Mar 31<sup>st</sup> | 17 | -17.9 | 0.14±0.011 a | 0.037±0.005 c | 0.103±0.007 ab | 76.471 | 20.19 |

(Means flanked with different letters at the same column are significantly different at $P<0.05$ )

**Fig 4.**
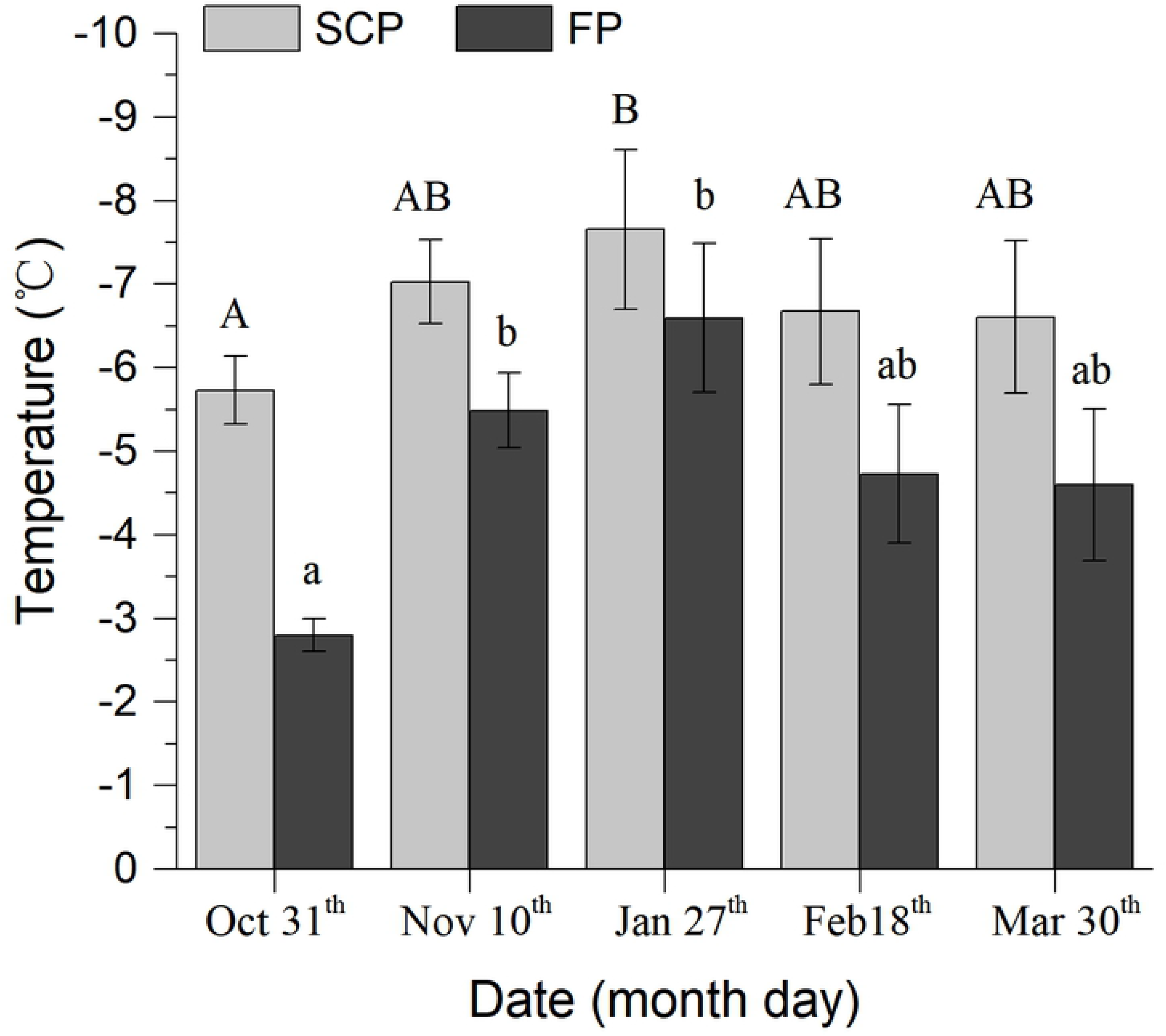
SCP and FP of *S. inferens* larvae during the overwinter period. Same pattern bar with different letters on top shows a significant difference at *p*< 0.05.

After a 30 minute SCP test, 91.667% of the tested *S. inferens* larvae were dead when returned to 24 °C immediately after freezing on October 31^st^. 15.789% died in the January 27^th^ SCP test, and 76.471% died in the March 31^st^ test. The number of dead larvae after the test was significantly negatively correlated with the daily air mean temperature on the sampling date (*r*=−0.915, *p* =0.029) (Table 1).

### 3.4 Current and potential distribution of *S. inferens*

The ALLT_50_ was −4.5 based on a nonlinear regression: *y*=(1.03054)^*x*^ (*y*: Mortality rate, *x*: Temperature (°C), *R*^2^=0.878, *p*<0.001, *n*=24) (Fig 5A). The ULLT_50_ was −9.713°C after being translated by a linear model between minimum daily air and underground temperature: *y*=0.6628\**x+*2.6267 (*y*: Underground, *x*: Air, *R*^2^=0.9161, *p*<0.0001, *n*=100) (Fig 5B). The regional survival ratio was calculated with both ALLT_50_ and ULLT_50_ based on low temperature threshold. ≥50% probability survival regions were below N 35° due to the ALLT_50_ and below N 40° due to the ULLT_50_ (Fig 6A).

**Fig 5.**
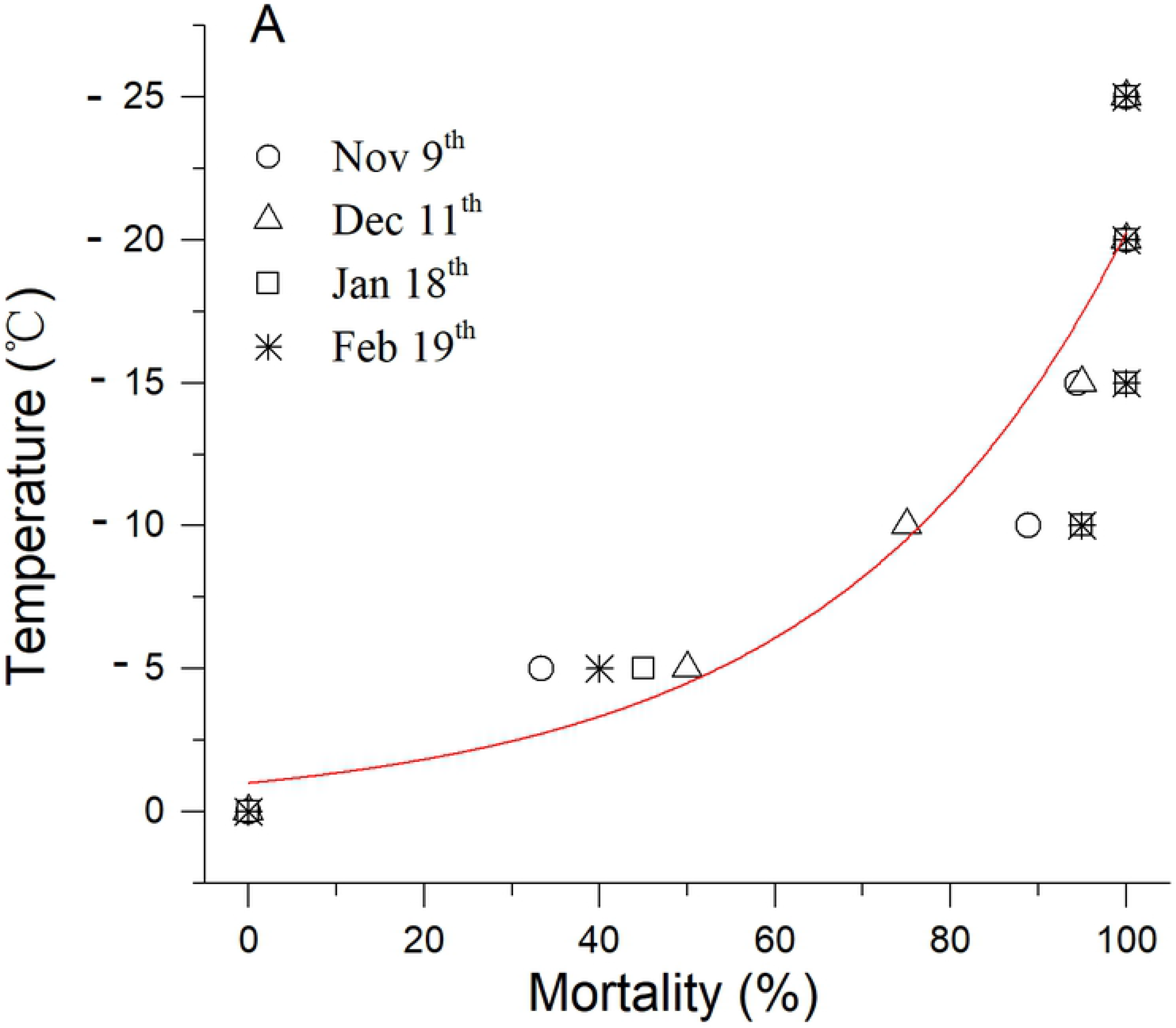
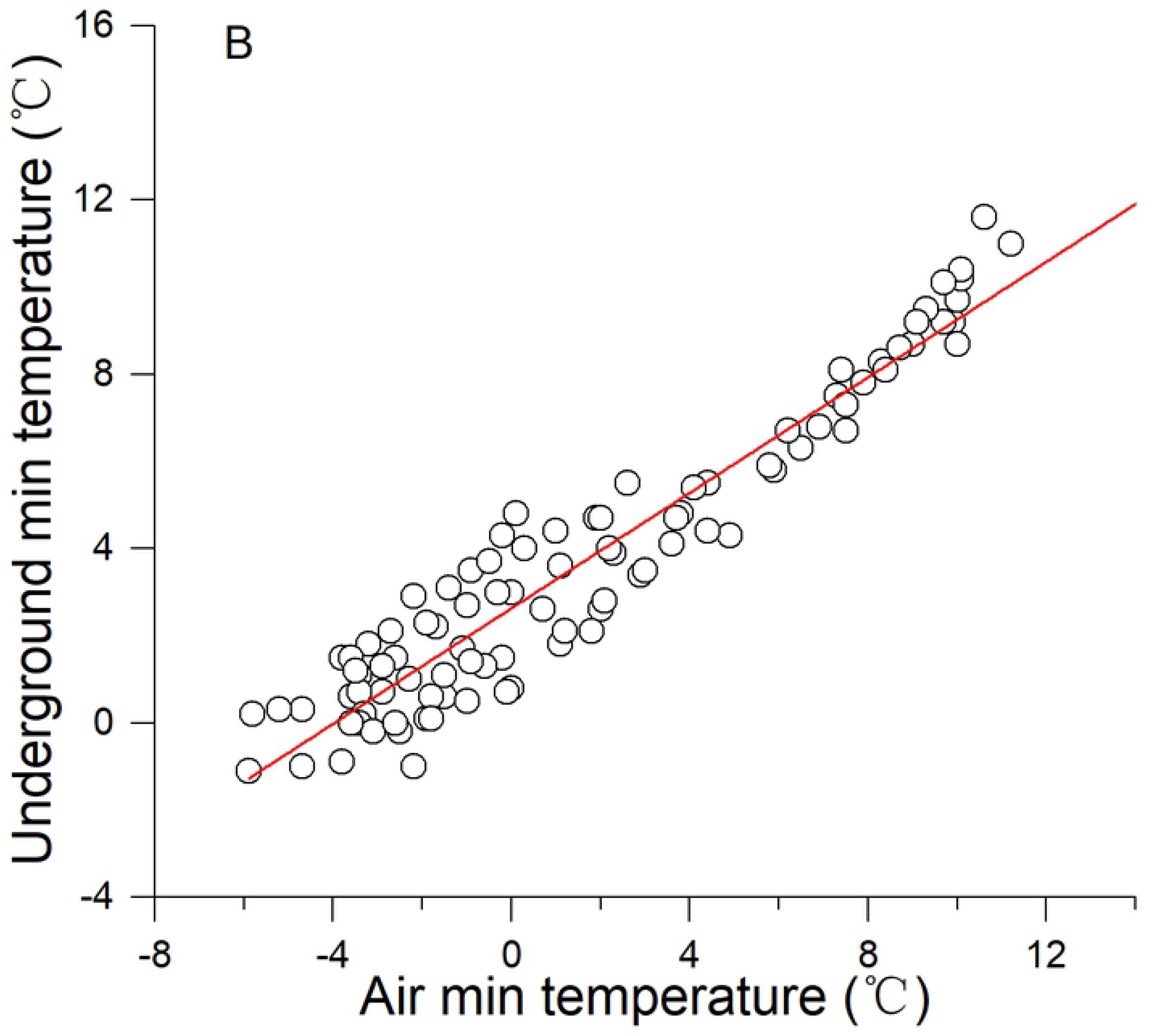
(A) Mortality rate of *S. inferens* larvae after lethal low temperature experiments and (B) the relationship between daily minimum air and UG temperature during the overwinter period.

**Fig 6.**
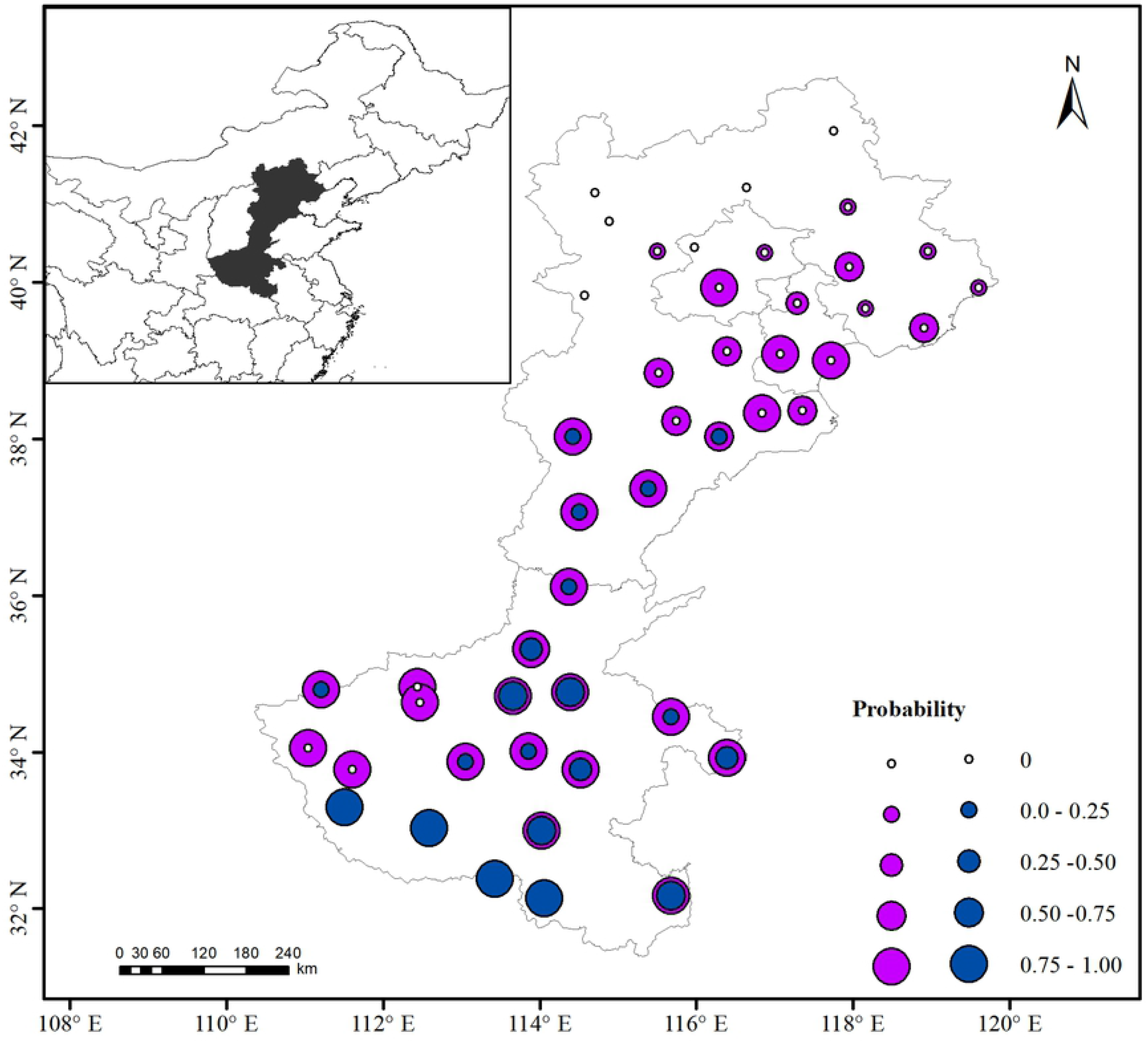
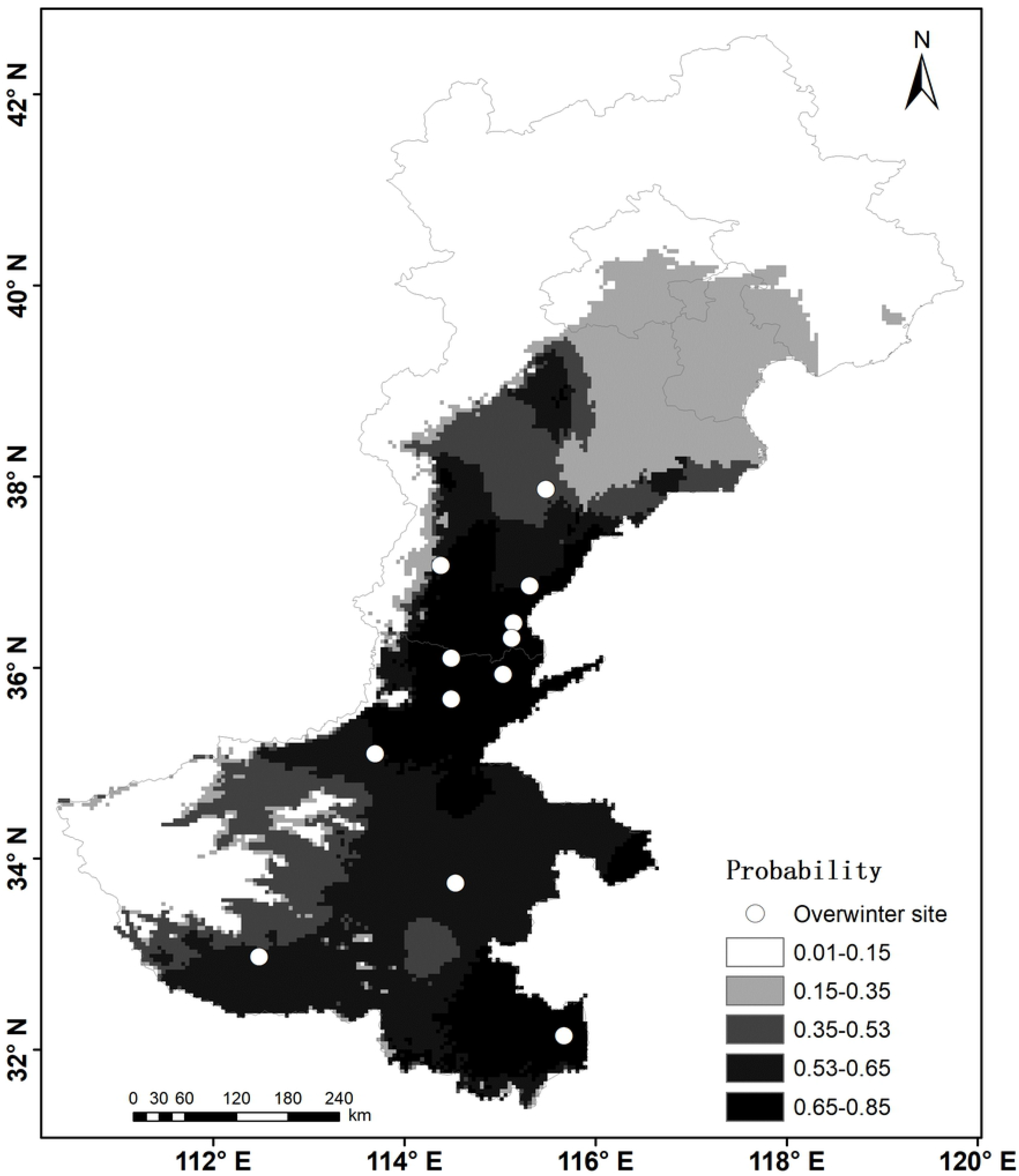
Current and potential distribution of *S. inferens* in northern China. (A) Potential distribution based on the low temperature threshold, the size of the circle in A means the survival probability, and the blue and purple circles are sourced from ALLT_50_ and ULLT_50_, respectively. (B) Current overwinter sites and predicted suitable habitat area modeling by MaxEnt, the darker color in B represents the better predicted habitat suitability. The white points are the new northern overwinter sites.

The MaxEnt model predicted potential distribution of *S. inferens* with a high accuracy test AUC value of 0.97 and training AUC value of 0.981(S1 Fig 2), and three temperature-variables: BIO6 (minimum temperature of coldest month), BIO10 (mean temperature of warmest quarter), and BIO2 (mean of monthly difference of maximum and minimum), which showed a top three permutation importance on the model (S1 Table 3). Model predictions closely matched the new overwinter site and also showed potentially suitable districts in north regions, and predicted suitable habitat areas of *S. inferens* below N 40°, and the most suitable areas were below N 38° (Fig 6B). This means that the underground overwinter temperature boundary defined by the ULLT_50_ is reliable, and overwinter strategies overcoming the winterkilling led to five latitude wider overwintering regions.

## 4 Discussion

There are many studies which show the latitudinal pattern of insects facing cold temperatures in winter. Higher latitude populations possess a stronger cold tolerance, a shorter chill-coma recovery time and a higher survival rate (Hoffmann et al.,2002; David et al., 2003; Chen et al., 2004; Rochefort et al.,2011; Tian et al., 2016). This research shows the overwintering location of the *S. inferens* population was placed closer to the ground with increasing latitude. In the new northern breeding area in winter, with the temperature decreasing, *S. inferens* larvae gradually climbed down and drilled into the base of the maize stalk under the soil surface before the coldest temperature arrived. Larvae remaining above ground in winter were all winterkilled. Compared to the year-round site at N 19° in the south (Zhou, 1985), it is an obvious behavioral evolution strategy of *S. inferens* to survive, to find the warmer microclimate of the overwintering site in the north in winter. Insects usually select warmer micro-habitats to survive. *Coccinella septempunctata* L., *Ceratomegilla undecimnotata* (Schneider), *Hippodamia variegata* (Goeze) *Harmonia axyridis* (Pallas) and *Coccinella septempunctata* select human houses, isolated grass tussocks or under covered stones as overwintering sites (Honek et al.,2007; Labrie et al., 2008; Koštál et al., 2014). The underground-forward behavior of *S. inferens* is also to avoid the chill damage in the northern area. Similarly, *Sesamia nonagrioide*, the same family but another species, also show underground-forward behavior in winter (Gillyboeuf et al., 1996).

Physiological adaptation of insects for low temperatures is by partial dehydration, increasing body fluid osmolality, and the accumulation of a complex mixture of winter specific metabolites, which strengthen their cold hardiness (Rozsypal et al., 2013; Koštál et al., 2014). In this paper, the mean SCP of larvae which were collected from the field was −7.653±0.952°C (lowest), and the mortality rate of the population was 15.789% (lowest) after a 30 minute cold hardiness test in January (middle winter). Meanwhile, the water contents gradually decreased and were lowest on Feb 18^th^ which shows the diapaused insects transfer fluidity into antifreeze proteins and glycoproteins (Rozsypal et al., 2014). When insects begin to recover in spring the fluidity increases. In this research, insects survived when body temperature was under freezing point and showed an extreme low temperature tolerance capability meaning that the population of *S. inferens* larvae in the northern breeding area was a moderately freeze-tolerant one. These results correspond with Guo et al. (2002) and Sun et al. (2014) where the pest hosts on rice. Cold hardiness is also related to the host insects have fed on (Morey et al., 2016; Alford et al., 2016) and the difference in the cold tolerance mechanisms between maize and rice need to be further investigated in the new breeding region.

Minimum temperatures in the coldest month are linearly related to insect mortality (Hoffmann et al., 2002). ALLT_50_ or the lethal temperature (LTe50) of insect cold exposure are the best predictors of cold distribution limits and are usually used to assess the potential distribution according to the annual minimum temperature (Ungerer et al., 1999; Andersen et al., 2015). This research used the same method to show a northern distribution considering the ALLT_50_ in locations under N 35°. However, depending on the ULLT_50_, the results showed a northerly distribution below N 40° when insects overwinter underground. This was also in agreement with the results of the MaxEnt modeling, which show below N 40° predicted suitable habitat areas. Distribution of insects is highly impacted by climatic factors (temperature, moisture, humidity and their variations), especially the effects of temperature (Aregbesola et al., 2019). MaxEnt integrates insect occurrence records with climatic and other environmental variables and the MaxEnt model of this study shows a similar result to de la Vega (2015), that the minimum temperature of the coldest month was the important abiotic factor restricting their geographic distribution. This research examined the geographical distributions of *Triatoma infestans* and *Rhodnius prolixus*. The MaxEnt model also included many other important parameters such as the precipitation, which showed a high contribution in the model. It is usually used to predict potential distributions of insect pests (Kumar et al., 2014). Although artificial because the linear model based on the point data might not represent the actual regional patterns, the linear relationship between air and soil temperature has indeed been studied in different areas (Parton and Logan, 1981; Horton, 2012), Hong (2014) has taken advantage of this linear relationship from 96 weather stations in Shanxi, China, and classified the overwintering sites of the southern root-knot nematode. These results also show that the underground low temperature threshold defined by the ULLT50 could be extrapolated to other underground overwinter species.

The population behavior of animals which escape cold weather usually by large-scale migration into a warmer habitat, will lead to relocated populations regionally, but not all individuals can successfully fly to suitable places. *S. inferens* have a weak flight tendency and capability in the field (Han et al., 2012). This paper shows another behavioral strategy of *S. inferens* larvae by locally searching for a warmer place by drilling into the ground in early winter. The microclimate of the overwintering site could result in the *S. inferens* colonizing more northerly latitudes and the insect’s physiological and biochemical changes may be a factor favoring this northern expansion leading to a higher winter survival chance. Past studies have shown that *S. inferens* are distributed below N 34° degrees north latitude. This study also documents a new location for *S. inferens*: a northern distribution record of N 40°. A large amount of research has shown that the overwintering boundary of many insects has been reportedly spreading poleward due to global warming (Wheatley et al., 2017). It cannot be overlooked that the warmer changes in northern China in these two centuries have driven the change of animal redistribution (Wu, 2016), but the strategy of behavioral or physiological changes in this research to overcome winter will certainly help to colonize a five degree more northerly latitude. For maize producers, ploughing the maize stubble in autumn after harvesting will lead to an increased mortality of *S. inferens* larvae due to cold exposure. This study allows us to propose a simple non-pollutant pest control method in northern China.

## Data Acessability

All data used in this manuscript is displayed in this context or provided as supporting information in the file: Supplementary Material 1.

## Author Contributions

JR and HQ conceived and designed the experiments. GP, HX and CB performed the overwinter investigation. JR analyzed the raw data. JR, GP, HX, CB, CH, QC, BH, HL, ZC, and HQ finalized the manuscript. All authors read and approved the final manuscript.

## Acknowledgements

We thank Chao Han and Mingyong Ma for carrying out the overwinter investigation on Nanjing and Changsha, respectively. We thanks those who contributed to discussions of the ideas presented in this paper. Instructive comments from the anonymous reviews will greatly improved this manuscript.

## Supporting information

**S1 Fig 1. The receiver operating characteristic (ROC) curve of the MaxEnt model.**

**S1 Fig 2. AUC of different environmental variables based on results of jackknife tests in the MaxEnt model.**

**S1 Table 1. The live number of *S. inferens* larvae at northern overwinter sites.**

**S1 Table 2. The worldwide distribution of *S. inferens***

**S1 Table 3. The permutation importance and relative contributions of the environmental variables in the MaxEnt model.**

